# Influence of perceptual load on attentional orienting in post-stroke fatigue: a study of auditory evoked potentials

**DOI:** 10.1101/2022.03.17.484808

**Authors:** William De Doncker, Annapoorna Kuppuswamy

## Abstract

**Objective:** Increasing perceptual load alters behavioural outcomes in post-stroke fatigue. While the effect of perceptual load on top-down attentional processing is known, here we investigate if increasing perceptual load modulates bottom-up attentional processing in a fatigue dependent manner.

**Methods:** In this cross-sectional observational study, in twenty-nine first-time non-depressed stroke survivors, an auditory oddball task consisting of target, standard and novel tones was performed in conditions of low and high perceptual load. Electroencephalography was used to measure auditory evoked potentials. Perceived effort was rated using the visual analog scale at regular intervals during the experiment. Fatigue was measured using the fatigue severity scale. The effect of fatigue and perceptual load on behaviour (response time, accuracy, and effort rating) and auditory evoked potentials (amplitude and latency) was examined using mixed model ANOVAS.

**Results:** Response time was prolonged with greater perceptual load and fatigue. There was no effect of load or fatigue on accuracy. Greater effort was reported with higher perceptual load both in high and low fatigue. p300a amplitude of auditory evoked potentials (AEP) for novel stimuli was attenuated in high fatigue with increasing load when compared to low fatigue. Latency of p300a was longer in low fatigue with increasing load when compared to high fatigue. There were no effects on p300b components, with smaller N100 in high load conditions.

**Interpretation:** High fatigue specific modulation of p300a component of AEP with increasing load is indicative of distractor driven alteration in orienting response, suggestive of compromise in bottom-up selective attention in post-stroke fatigue.

## Introduction

Post-stroke fatigue (PSF), a feeling of extreme exhaustion, is a commonly reported symptom with a major impact on quality of life and mortality^1–4^. Despite high prevalence of fatigue after stroke, the underlying neural pathophysiology is poorly understood.

Heightened effort perception has been proposed as a cause of fatigue^5,6^, with cognitive dysfunction underpinning altered effort perception. Tests of selective attention such as trail making tests and flanker tasks, take longer to complete in those with post-stroke fatigue^7–10^. An in-depth analysis of the driving factors of poor performance in selective attention tasks indicate a problem with stimulus encoding and/or action execution rather than longer decision time^10^. Recent results from our lab suggest stimulus encoding, specifically distractor stimuli, are linked to high levels of fatigue^11^. We observed that with greater task related cognitive load (working memory demand), poorer distractor suppression was seen in high fatigue. Such poor distractor suppression increases perceptual load, but did not influence representation of top-down attentional set (anticipated target stimuli). However, its effect on processing of unanticipated novel stimuli (bottom up attention) remains unknown. Greater perceptual load has previously been linked to diminished bottom-up processing both in healthy humans^12^, and in those with neurological disorders^13^ indicative of reduced capacity to process unexpected sensory stimuli. In post-stroke fatigue, with poor distractor suppression resulting in greater perceptual load, an attenuated bottom-up attentional response is anticipated.

Bottom-up attention is investigated using a three-tone auditory oddball paradigm which evokes event-related potentials (ERPs) recorded using electroencephalography (EEG). The fundamental premise of this paradigm is that when one is tasked with attending to a target tone presented amongst non-target tones, a third, non-target novel tone when presented, despite not requiring a response, involuntarily elicits a response, commonly named an orienting response, indicative of bottom-up processing. Such a response is reflected in the latency and amplitude components of p300a element of ERP, with other components reflecting other attentional and perceptual processes^14^. The effect of perceptual load on the orienting response will be investigated by introducing a ‘noise’ condition where the task is performed in the presence of background noise, thereby increasing the perceptual load of the task.

In summary, here we investigate if self-reported fatigue levels in chronic stroke survivors is associated with diminished orienting response driven by perceptual load in a three-tone auditory oddball task.

## Material and Methods

### Participants

This was a cross-sectional observational study approved by the London Bromley Research Ethics Committee (REC reference number: 16/LO/0714). Twenty nine stroke survivors were recruited via the Clinical Research Network from the University College NHS Trust Hospital (UCLH), a departmental Stroke Database and from the community. All stroke survivors were screened prior to the study based on the following criteria: first-time ischaemic or haemorrhagic stroke; stroke occurred at least 3 months prior to the study; no clinical diagnosis of any other neurological disorder; physically well recovered following their stroke defined as grip strength and manual dexterity of the affected hand being at least 60% of the unaffected hand assessed using a hand-held dynamometer and the nine-hole peg test (NHPT) respectively; not taking anti-depressants or any other medication that has a direct effect on the central nervous system; not clinically depressed with depression scores ≤ 11 assessed using the Hospital Anxiety and Depression Scale (HADS). ^15^. Following written informed consent in accordance with the Declaration of Helsinki, twenty-nine stroke survivors participated in the study (Table 1).

**Table 1.**
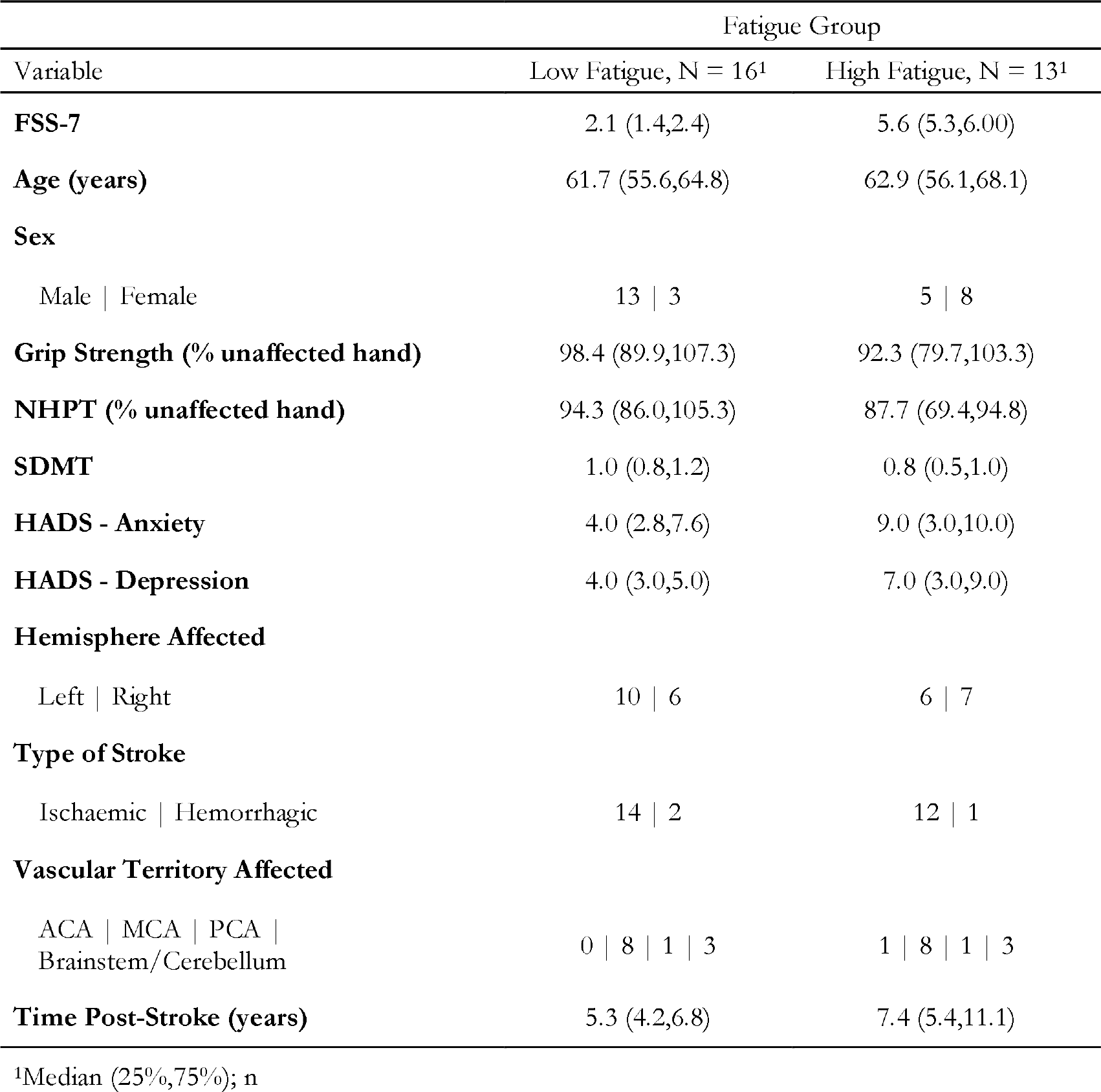
Demographics of the high and low fatigue groups that took part in the study. The median and interquartile range is shown for continuous variables, while the count is shown for categorical variables. NHPT = Nine Hole Peg Test, SDMT = Symbol Digit Modalities Test, HADS = Hospital Anxiety and Depression Scale, ACA = Anterior Cerebral Artery, MCA = Middle Cerebral Artery, PCA = Posterior Cerebral Artery.

### Fatigue

Two measures of fatigue were captured at the start of the study, trait and state fatigue. Trait fatigue represents the experience and impact of fatigue on day to day living for a pre-determined time leading up to the day of testing, whereas state fatigue characterizes fatigue at a given moment in time. Trait fatigue was quantified using the FSS-7, a seven-item questionnaire that rates fatigue over a 1 week period preceding the date of testing^16^. State fatigue was quantified using a visual analogue scale (VAS) ranging from zero to ten in steps of 1 (Not at all tired to extremely tired).

### Stimuli and Procedure

Participants were seated 70 centimeters from the monitor (Dell U240 24-inch monitor, at a screen resolution of 1280x768) and made their responses using a standard USB keyboard. The experiment was controlled using the Psychophysics Toolbox for Matlab^17,18^, running on a Windows computer. A three-stimulus auditory oddball paradigm was used to elicit N100, P300a and P300b event-related potentials (ERPs) both in the presence and absence of noise (Figure 1). Stimuli consisted of 1,200 binaural, 80 dB tones of 150 ms duration presented to the participants through headphones (Sennheiser, HD 569), divided into twelve blocks (100 stimuli per block) lasting approximately four minutes each. In 50% of the blocks (6 of 12), participants performed the oddball task with no noise, while in the remaining 50% of the blocks, participants performed the oddball task in the presence of noise. The ‘noise’ was an ecologically valid recording of chatter in the café (babble) played at 65 dB. The order of block presentation was counterbalanced across participants. Twelve percent of the stimuli (144 in total, 12 per block) were target tones (1.5 kHz tone), 12% of the stimuli (144 in total, 12 per block) were “novel” sounds (a cricket sound, a sneeze etc), and 76% were standard tones (1 kHz tone), with an inter-stimulus interval varying between 1.8 and 2.2 seconds. Participants were instructed to press a button on the keyboard with the index finger of their right hand in response to target tones only, while keeping their eyes on a fixation cross. This allowed us to have a measure of response time (time from auditory stimulus onset to button press) and accuracy (% correct of button press) for each participant. Participants were also asked to rate the effort required to complete a block of the task, on a VAS ranging from zero (very easy) to ten (very hard). Before the start of the experiment participants completed practice trials (15 correct trials), both in the presence and absence of background noise, to ensure they familiarized themselves with the task demands.

**Figure 1.**
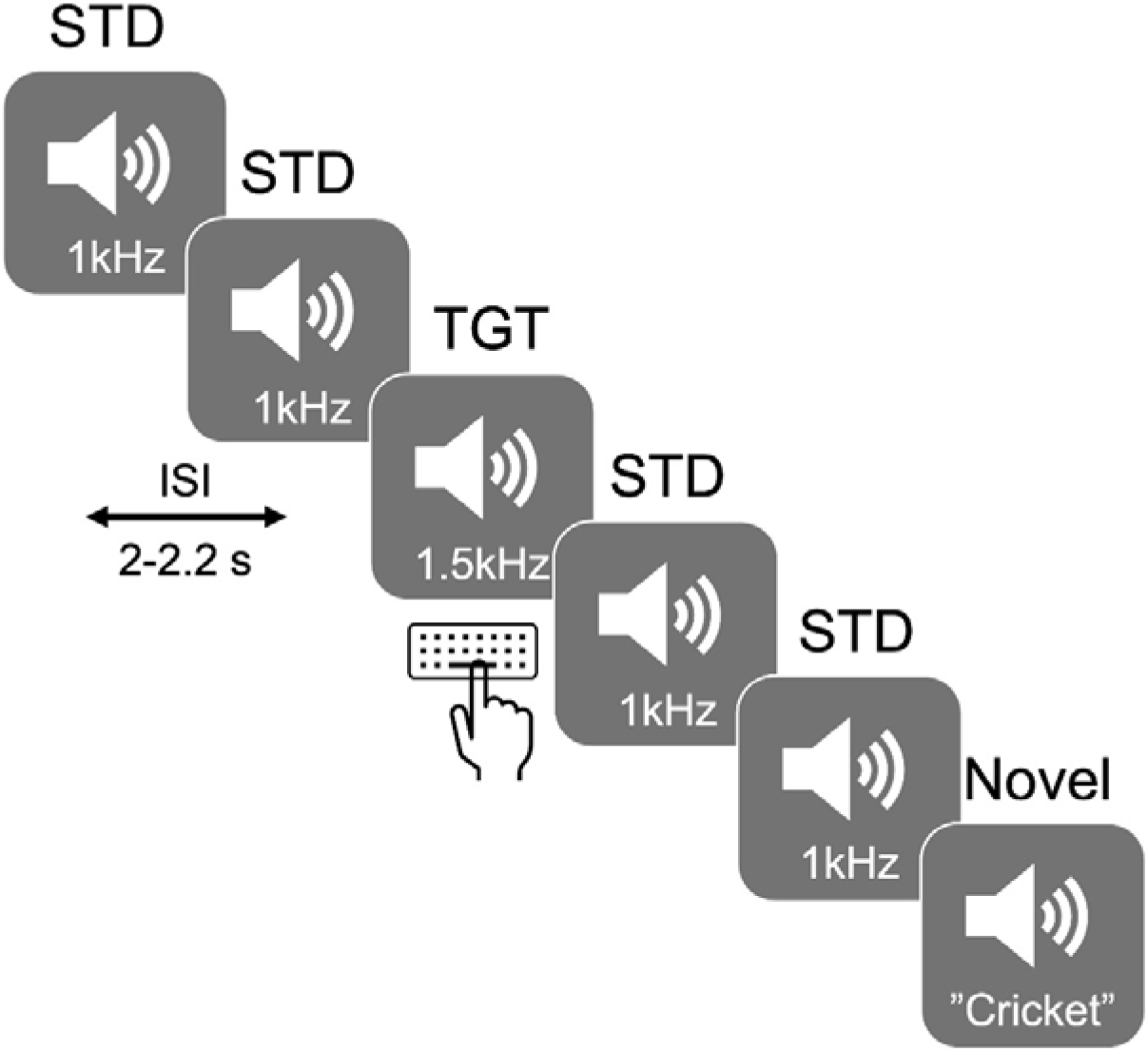
Illustration of the task design. Target (TGT) tones at 1.5 kHz requiring a requiring a response via a keyboard and Novel tones were presented amongst a sequence of standard (STD) tones at 1 kHz.

### EEG Acquisition

Whole-scalp electroencephalography (EEG) data was recorded using a 64-channel cap array (ActiCap, Herrsching, Germany) and a BrainAmp EEG amplifier system (BrainProducts, Gilching, Germany). The 64 electrodes were positioned on the cap in accordance with the 10-20 international EEG electrode array. During online recordings, channels FCz and AFz were used as the reference and ground respectively. Impedances were kept below 10 kΩ throughout the recording. The EEG signal was sampled at 1 kHz and visualized online using the BrainVision Recorder Software (BrainVision Recorder, Version 1.21.0102 Brain Products GmbH, Gilching, Germany). Event markers were sent from the stimulus presentation PC to the BrainAmp amplifier via the TriggerBox which allows one to send accurate triggers via a USB port with millisecond precision.

### EEG Analysis

EEG data were pre-processed and analysed offline using EEGLAB^19^, ERPLAB^20^ and customized MATLAB scripts (Mathworks, Inc., MA, USA). EEG data was down-sampled to 250 Hz and subsequently band-pass filtered between 0.5 and 30 Hz with a zero phase-shift IIR Butterworth filter (24 dB/Oct). Noisy channels were identified and removed using automated procedures. To identify and remove ocular movements and blink artefacts from the EEG data, an independent component analysis (ICA) implemented within EEGLAB was used. The components were visually inspected and those containing ocular movements or blink artifacts were removed. The previously removed channels were then interpolated back into the dataset and finally, the EEG data was re-referenced against the grand average of all scalp electrodes.

### ERP

The pre-processed EEG data was segmented into epochs of -200 ms to 800 ms time locked to the auditory stimulus onset and baseline corrected using the 200 ms pre-stimulus period. Individual epochs were inspected using a 200 ms sliding time window in steps of 100 ms across the entire length of the epoch for voltages exceeding ± 100 µV. These trials were subsequently excluded from the analysis (2.5 % ± 3.4) of trials. All artefact-free epochs were then averaged for each of the three conditions (Standard, Target, Novel) and filtered using a low-pass IIR Butterworth filter of 12 Hz. The mean number of trials remaining was comparable between groups for each condition (Table 2).

**Table 2.** Number of trials for each condition across the two fatigue groups. The mean number of trials and the standard deviation across each condition, as well as the p-value for the difference between the two groups is shown.

| Tone Type | Fatigue Group |  |  |
| --- | --- | --- | --- |
|  | Low, N = 16<sup>1</sup> | High, N = 13<sup>1</sup> | p-value<sup>2</sup> |
| **Standard** | 445.94 (14.06) | 443.62 (15.76) | 0.6 |
| **Standard with Noise** | 439.75 (25.40) | 444.92 (23.62) | 0.9 |
| **Target** | 70.44 (1.90) | 70.23 (3.03) | 0.8 |
| **Target with Noise** | 67.69 (6.39) | 70.92 (2.10) | 0.093 |
| **Novel** | 71.06 (1.34) | 70.15 (2.85) | 0.7 |
| **Novel with Noise** | 69.56 (4.35) | 70.69 (2.69) | 0.9 |

<sup>1</sup>Mean (SD)
<sup>2</sup>Wilcoxon rank sum test

The mismatch negativity (MMN) wave was estimated by subtracting the grand average of the standard tones from the grand average of the novel tones (MMNa) and from the grand average of the target tones (MMNb) in each participant. The N100 amplitude was defined as the instantaneous peak negative amplitude between 50 and 200 ms from the auditory stimulus onset at electrode Cz across the three conditions; the P300a amplitude was defined as the instantaneous peak positive amplitude between 250 and 450 ms from the auditory stimulus onset at electrode CPz in MMNa wave; the P300b amplitude was defined as the instantaneous peak positive amplitude between 280 and 650 ms from the auditory stimulus onset at electrode Pz in the MMNb wave. The latency of each peak across the three ERPs was also recorded for statistical analysis. These three midline locations were chosen as ERP responses were largest at these electrode sites.

### Statistical Analysis

All statistical analyses were performed using R (RStudio Version 1.2.5033). Spearman rank correlations were used to identify the association between trait fatigue (FSS-7) and all continuous demographic measures (age, grip strength, NHPT, HADS-Depression, HADS-Anxiety and Time Post-Stroke), while Wilcoxon rank sum tests were used to identify the association between trait fatigue and all categorical demographic measures (sex, hemisphere affected, type of stroke and vascular territory affected).

The distribution of the dependent variable was assessed using the Shapiro-Wilk’s test of normality, while homogeneity of variances was assessed using the Levene’s test. As all data was normally distributed, group differences in the behavioural (response time, accuracy, and effort) and ERP data (amplitude and latency) were examined using a mixed analysis of variance with group (Low Fatigue and High Fatigue) as the between subject factor and noise (Off/On) as the within subject factor. Differences were considered statistically significant at the level of p < 0.05. Generalised eta squared (η^2^) was reported for the group effect sizes. Bonferroni corrected pairwise t-tests were used to assess simple main effects in the post-hoc analysis.

## Results

### Participant Demographics

Twenty-nine stroke survivors completed the study (11 females and 18 males). The median FSS-7 score was 5.29 (IQR = 2.57) in females and 2.50 (IQR = 2.46) in males. The Wilcoxon test showed that the difference in FSS-7 score was marginally non-significant (p = 0.05, effect size = 0.37). Spearman rank correlations between trait fatigue (FSS-7) and all continuous demographic measures revealed a significant positive association between trait fatigue and HADS-Depression (Spearman ρ = 0.41, p = 0.03), while no other variable correlated with trait fatigue. The median FSS-7 score in those with right and left hemisphere strokes was 4.43 (IQR = 3.00) and 2.71 (IQR = 3.75) respectively (Wilcoxon test: p = 0.50, effect size = 0.13). The median FSS-7 score in ischemic strokes was 2.93 (IQR = 3.46) and 3.86 (IQR = 2.29) in hemorrhagic strokes (Wilcoxon test: p = 0.51, effect size r = 0.13). The median FSS-7 score in ACA strokes was 6.85, in MCA strokes was 3.36 (IQR = 3.25), in PCA strokes was 3.07 (IQR = 2.07), and 4.21 (IQR = 2.68) in Brainstem/Cerebellum strokes (Kruskal-Wallis test: p = 0.57, η^2^ = -0.04). A spearman rank correlation between FSS-7 and the Time Post-Stroke showed no significant association (Spearman ρ = 0.08, p = 0.67). Any meaningful interpretation of the effect of the type of stroke and vascular territory affected on FSS-7 in the current cohort of stroke survivors is difficult given the skewed numbers.

### Behaviour

The response time (RT) results are shown in Figure 2A. As the RT data was not normally distributed, the data were log transformed and entered into a mixed ANOVA with noise (Off/On) as the within-subject factor and fatigue (Low Fatigue/High Fatigue) as the between subject factor. There was a main effect of noise (F_(1,27)_ = 8.12, p = 0.01, η^2^ = 0.01) and a main effect of fatigue (F_(1,27)_ = 5.40, p = 0.03, η^2^ = 0.16) but no interaction between noise and fatigue (F_(1,27)_ = 0.17, p = 0.69, η^2^ < 0.01). Pairwise t-tests showed a significant difference in RT between the two fatigue groups in the noise off (p = 0.025) and the noise on (p = 0.036) conditions.

**Figure 2.**
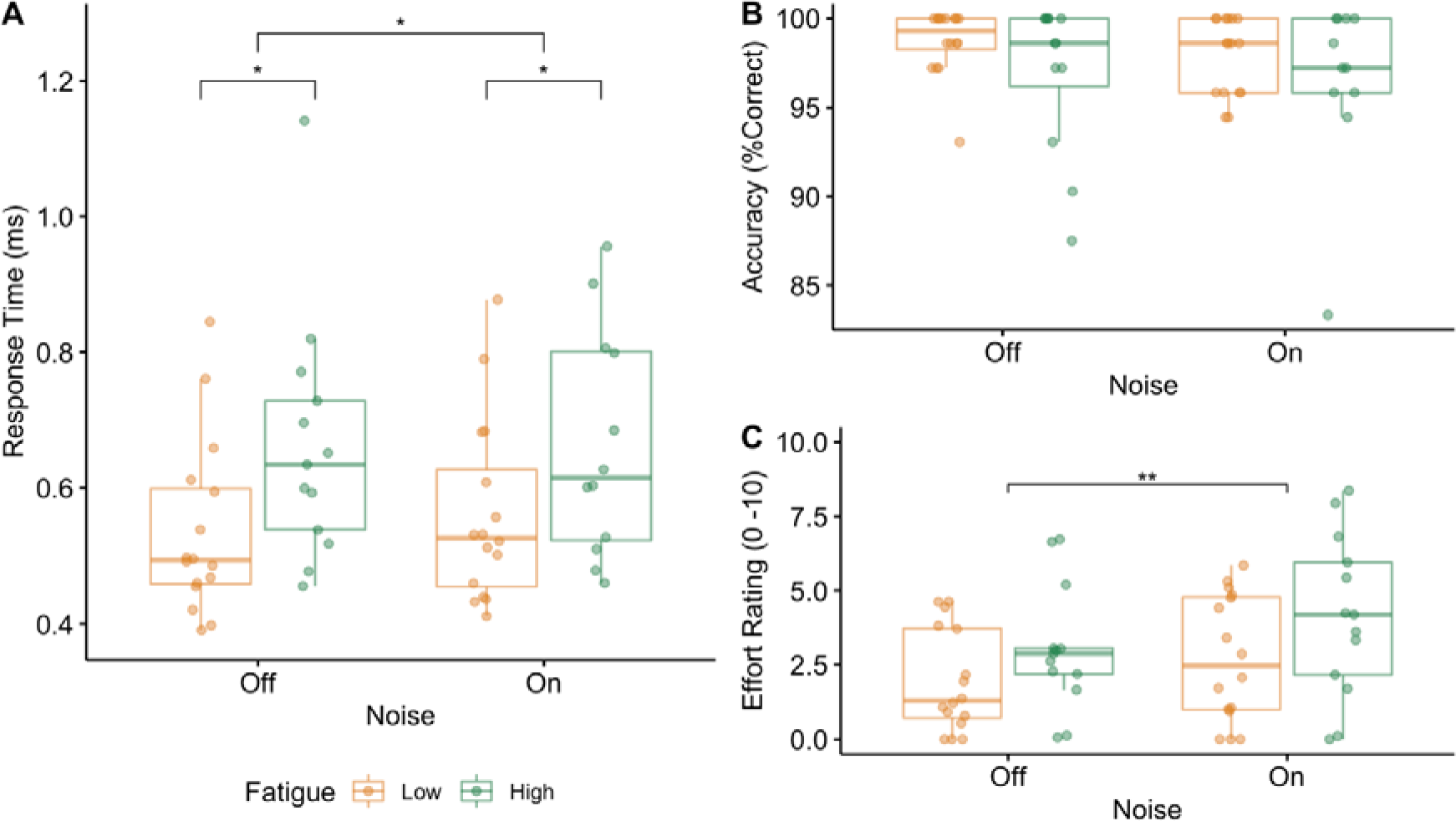
Behavioural results. Box plots with individual data points for the low fatigue group in yellow and the high fatigue group in green for response time (A), accuracy (B) and effort rating (C) across the two background noise conditions (Off and On). Significant differences between the groups are indicated with asterisks (*p < 0.05).

The accuracy results are shown in Figure 2B. The mixed ANOVA revealed no effect of noise (F_(1,27)_ = 1.21, p = 0.28, η^2^ = 0.01), no effect of fatigue (F_(1,27)_ = 3.36, p = 0.08, η^2^ = 0.10) and no interaction between noise and fatigue (F_(1,27)_ = 0.45, p = 0.51, η^2^ < 0.01).

The effort rating results are shown in Figure 2C. The mixed ANOVA revealed a main effect of noise (F_(1,27)_ = 10.50, p < 0.01, η^2^ = 0.05), no effect of fatigue (F_(1,27)_ = 2.80, p = 0.11, η^2^ = 0.08) and no interaction between noise and fatigue (F_(1,27)_ = 0.37, p = 0.55, η^2^ < 0.01).

### P300a amplitude and latency

Grand average waveforms and topographic maps for P300a are shown in Figure 3. The mixed ANOVA with the amplitude of the P300a as the dependent variable revealed a main effect of noise (F_(1,27)_ = 37.18, p < 0.01, η^2^ = 0.12), no effect of fatigue (F_(1,27)_ = 3.36, p = 0.08, η^2^ = 0.10) and a significant interaction between noise and fatigue (F_(1,27)_ = 8.50, p < 0.01, η^2^ = 0.03). Pairwise t-test showed that there was a significant difference in the amplitude of the P300a between the two fatigue groups in the noise on (p = 0.02) but not in the noise off (p = 0.45) conditions (Figure 4A). To further examine how the presence of noise differentially modulates the amplitude of the P300a across the two fatigue groups, the change in amplitude between the noise off and on conditions was calculated and compared between the two fatigue groups using a t-test (t-statistic = -2.75, df = 17.8, p = 0.01; Figure 4B).

**Figure 3.**
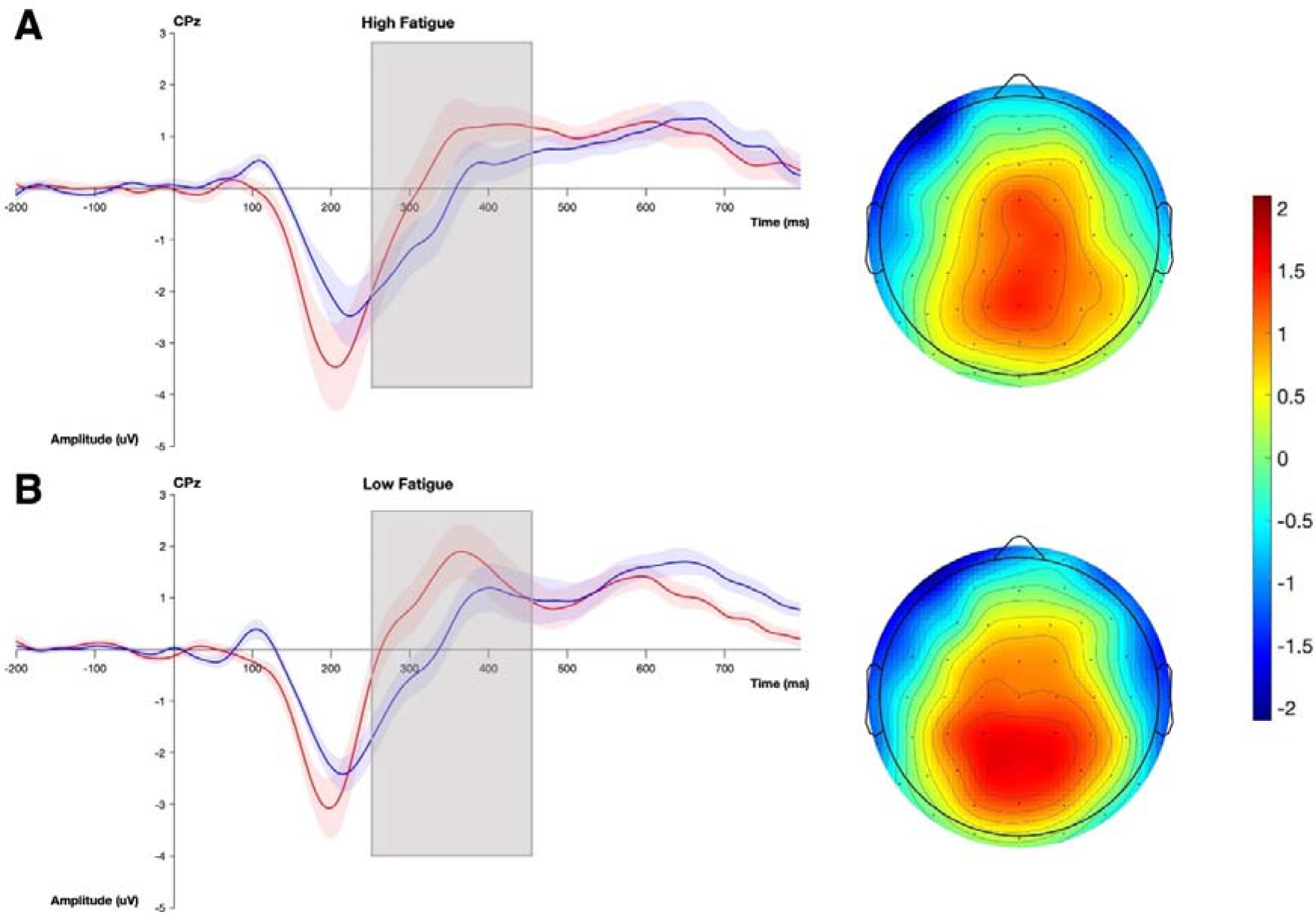
Grand average ERP of the P300a at electrode CPz and topographical plot for the high (A) and low (B) fatigue groups. The noise off condition is in red and the noise on condition is in blue. The topographical plots illustrate the difference between the two noise conditions. The grey box indicates the time window within which the P300a was measured

**Figure 4.**
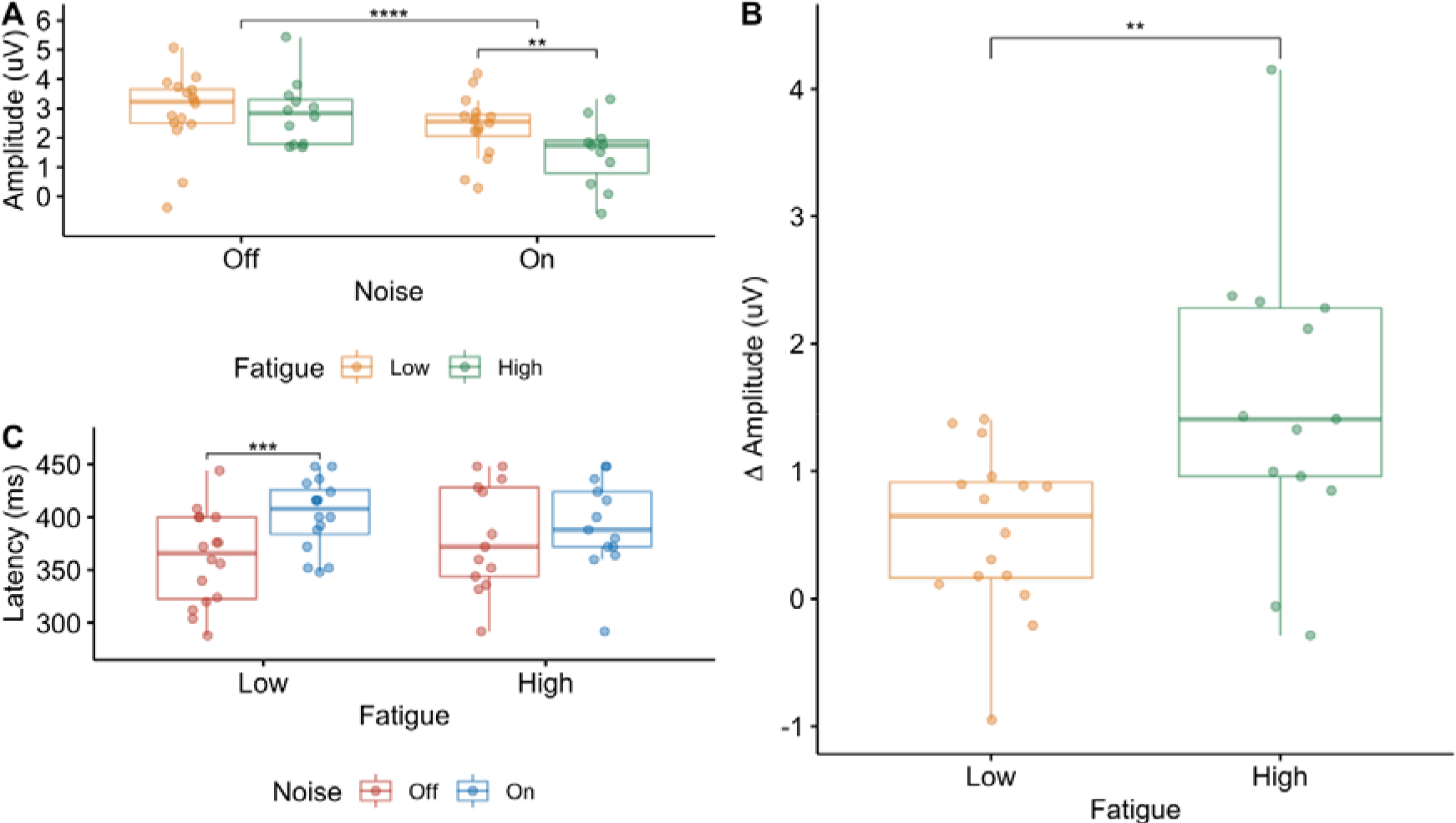
P300a results. Box plots with individual data points for the low fatigue group in yellow and the high fatigue group in green for the amplitude (A) and the difference in amplitude across the two noise conditions (B) for each fatigue group of the P300a. Box plots with individual data points for the latency of the P300a across the different noise conditions, with noise off in red and noise on in blue, is shown in panel C. Significant differences between the groups are indicated with asterisks (*p < 0.05).

The mixed ANOVA with the latency of P300a as the dependent variable (Figure 4C) revealed a main effect of noise (F_(1,27)_ = 13.80, p < 0.01, η^2^ = 0.09), no effect of fatigue (F_(1,27)_ = 0.12, p = 0.74, η^2^ < 0.01) and a significant interaction between noise and fatigue (F_(1,27)_ = 4.59, p = 0.04, η^2^ = 0.03). Pairwise t-tests showed a significant difference in the latency of the P300a between the two noise conditions in the low fatigue group (t-statistic = -4.87, df = 15, p < 0.01) but not in the high fatigue group (t-statistic = -0.95, df = 12, p = 0.36).

### P300b amplitude and latency

The mixed ANOVA with the amplitude of P300b as the dependent variable (Figure 5A) revealed no effect of noise (F_(1,27)_ = 0.03, p = 0.85, η^2^ < 0.01), no effect of fatigue (F_(1,27)_ = 0.002, p = 0.97, η^2^ < 0.01) and no significant interaction between noise and fatigue (F_(1,27)_ = 0.00006, p = 0.99, η^2^ < 0.01). The mixed ANOVA with the latency of the P300b as the dependent variable (Figure 5B) revealed no effect of noise (F_(1,27)_ = 0.03, p = 0.88, η^2^ < 0.01), no effect of fatigue (F_(1,27)_ = 0.53, p = 0.47, η^2^ = 0.012) and no significant interaction between noise and fatigue (F_(1,27)_ = 0.04, p = 0.84, η^2^ < 0.01).

**Figure 5.**
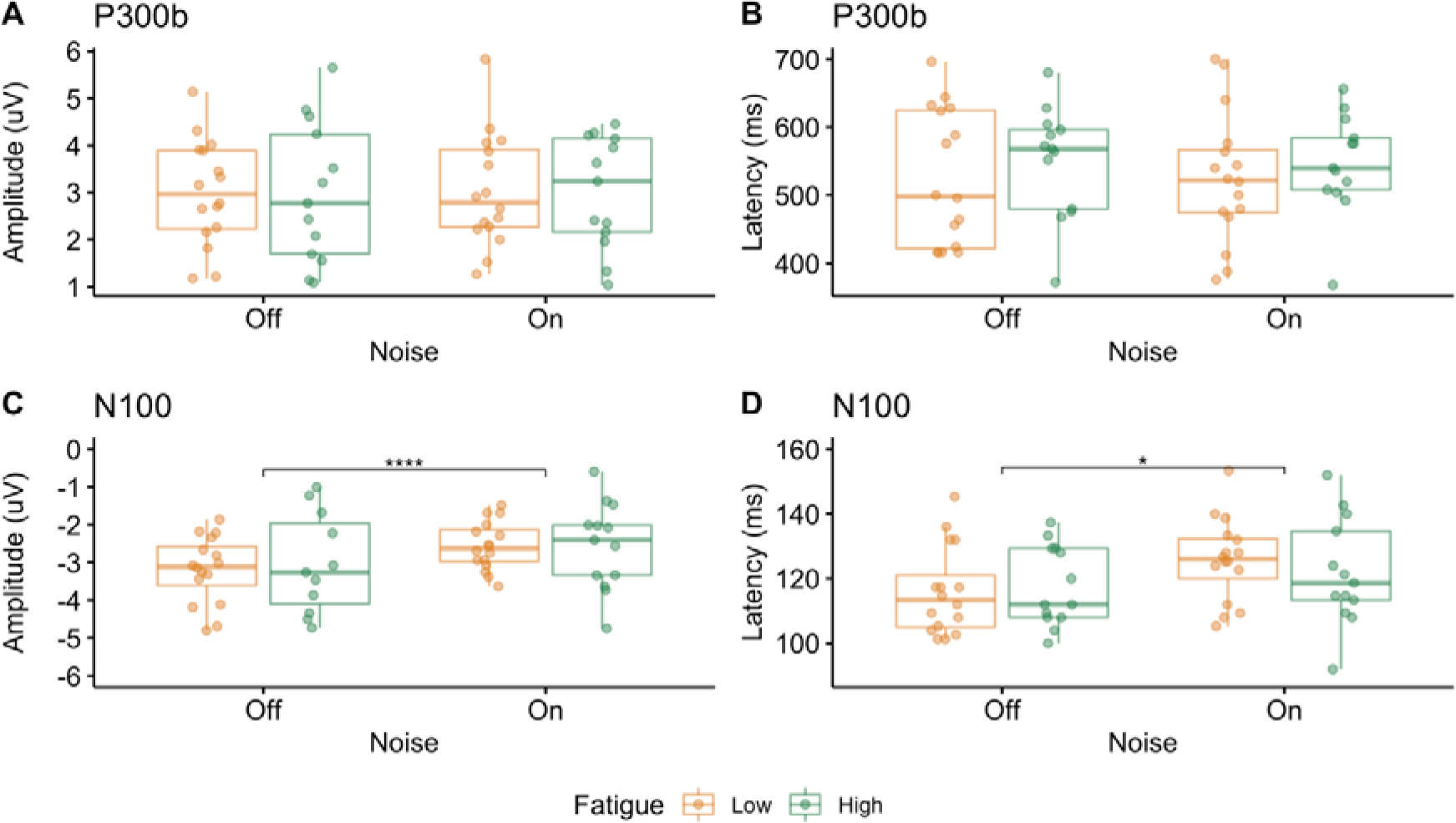
P300b and N100. Box plots with individual data points for the low fatigue group in yellow and the high fatigue group in green for the amplitude (A) and latency (B) of the P300b, as well as the amplitude (C) and latency (D) of the N100. Significant differences between the groups are indicated with asterisks (*p < 0.05).

### N100 amplitude and latency

The mixed ANOVA with the amplitude of the N100 as the dependent variable (Figure 5C) had an additional within-subject factor of Condition (Standard, Target, Novel) to examine whether the amplitude of the N100 was differentially modulated across the different conditions. The mixed ANOVA revealed no effect of condition (F_(2,54)_ = 0.69, p = 0.44, η^2^ = 0.01). The amplitude of the N100 was therefore averaged across the three conditions, and a mixed ANOVA was re-computed with noise as the only within-subject factor. The mixed ANOVA revealed a main effect of noise (F_(1,27)_ = 42.06, p < 0.01, η^2^ = 0.12), no effect of fatigue (F_(1,27)_ = 0.14, p = 0.71, η^2^ < 0.01) and no significant interaction between noise and fatigue (F_(1,27)_ =1.17, p = 0.21, η^2^ < 0.01). The mixed ANOVA with the latency of the N100 as the dependent variable (Figure 5D) revealed an effect of noise (F_(1,27)_ = 5.28, p = 0.03, η^2^ = 0.06), no effect of fatigue (F_(1,27)_ = 0.06, p = 0.81, η^2^ < 0.01) and no significant interaction between noise and fatigue (F_(1,27)_ = 0.83, p = 0.37, η^2^ = 0.01).

## Discussion

In twenty-nine minimally impaired, non-depressed, chronic stroke survivors, we show that fatigue severity scale score is predictive of a greater reduction in orienting response (P300a) with increasing perceptual load. An inverse relationship between fatigue and change in latency of orienting response was seen, with increase in perceptual load associated with longer latency of response in low fatigue compared to high fatigue. No effect of load or fatigue was observed on P300b while amplitude of N100 was reduced with increasing load but not fatigue. We also show that perceptual load prolongs behavioural response times irrespective of fatigue levels, and higher fatigue is associated with slower response times irrespective of perceptual load. Accuracy was unaffected by perceptual load and fatigue, however effort required to perform the task, as indicated by self-report, was higher with greater perceptual load and not affected by fatigue.

The main finding of this study is a fatigue dependent amplitude and latency modulation of P300a response with increase in task-irrelevant perceptual load. A lack of baseline difference in the orienting response between high and low fatigue shows that bottom-up processing is similar across groups in the absence of ‘noise’. However, increasing perceptual load by task irrelevant background noise results in a significant reduction in the orienting response only in high fatigue and not in low fatigue, highlighting the importance of perceptual load in the experience of fatigue. While this study did not directly measure noise encoding, our previous results demonstrating poor distractor suppression with increasing perceptual load suggests that noise related alteration in orienting response is likely driven by poor noise (distractor) suppression^11^. In healthy humans, fatigue inducing paradigms also result in poor orienting response only when the paradigm inducing fatigue has high perceptual load with low perceptual load paradigms having no effect on orienting response^12^. In Parkinson’s disease and traumatic brain injury, fatigue is associated with attenuated p300a while not affecting p300b components^13,21^, indicating similar neural processes might underlie fatigue across different neurological disorders. In disorders such as autism spectrum disorder and schizophrenia, changes in attentional responses, specifically bottom-up processing deficits manifest as a reduced capacity to meaningfully engage with the environment^22,23^. Individuals with fatigue also severely restrict their interaction with the environment, and what was previously attributed to reduced motivation due to fatigue, might be driven by abnormal sensory processing.

With increasing perceptual load, the latency of p300a lengthened in low fatigue but not in high fatigue. Latency of p300a reflects stimulus evaluation and attention allocation time^14^, and with increasing perceptual load one expects a lengthening of latency as seen in low fatigue. However, in high fatigue there was no effect of load on latency. On closer examination, we see that average latency in low load condition is similar to that of high load condition in low fatigue suggesting that in high fatigue irrespective of load it takes longer to evaluate stimuli and allocate attention. Fatigue in other diseases such as Parkinson’s disease^13^ and multiple sclerosis^24^ are also related to longer p300a latency, further highlighting the commonalities of fatigue across disorders.

There were no load specific behavioural consequences that were exclusive to the high fatigue group as would be expected from the load specific attenuation of orienting response in the high fatigue group. Both speed and accuracy of response, and task related effort were heightened with increasing perceptual load across both groups. A lack of behavioural effect could either be a result of the task difficulty not reaching a threshold for the underlying neural processing changes to reflect on behaviour, or, in keeping with previous results^25,26^, such altered sensory processing exclusively propagates to ‘sensory awareness’ networks and not to motor output pathways.

In summary, we show that processing of novel stimuli in the presence of background noise is significantly altered in PSF, indicating a problem with attentional orienting processes, one that selectively contributes to the feeling of fatigue without affecting performance. This evidence provides a physiological basis for self-reported fatigue-related difficulties of processing several streams of sensory information. These results, taken together with previous findings both in stroke and other disorders where compromise in dopamine dependent sensory processing is seen, paves the way for future research studies to investigate common fatigue mechanisms across diseases and develop new therapeutic interventions.

## Acknowledgements

We would like to acknowledge Paul Hammond for all the technical help and assistance throughout this study, as well as Dr Chi-Hsu Wu for his help in collecting part of the data.

## Author Contributions

Conception and design (AK, WDD); acquisition and analysis of data (WDD); drafting a significant portion of the manuscript or figures (WDD).

